# Loss of Wdr23 drives neuronal mitochondrial biogenesis

**DOI:** 10.1101/2023.09.05.556455

**Authors:** Ronald Irwin, Jiahui Liu, Chatrawee Duangjan, Sean P. Curran

**Author notes:** equal contribution.

## Abstract

Mitochondrial adaptation is important for stress resistance throughout life. Here we show that WDR23 loss results in an enrichment for genes regulated by nuclear respiratory factor 1 (NRF1), which coordinates mitochondrial biogenesis and respiratory functions, and an increased steady state level of nuclear coded mitochondrial resident proteins in the brain. *Wdr23KO* also increases the endogenous levels of insulin degrading enzyme (IDE) and the relaxin-3 peptide (RLN3), both of which mediate mitochondrial metabolic and oxidative stress responses. Taken together, these studies reveal an important role for WDR23 as a component of the mitochondrial homeostat in the murine brain.

**HIGHLIGHTS:**

- Loss of Wdr23 increases nuclear-coded mitochondrial resident proteins.
- Promoters of transcripts dysregulated in the hippocampus of *Wdr23KO* mice are enriched for NRF1 regulatory sequences.
- Insulin degrading enzyme (IDE) expression, which can localize to the mitochondria, is increased in the brain tissues lacking WDR23.
- *Wdr23KO* animals have increased expression of relaxin-3 (RLN3) peptide, but not RLFP3 receptor.

## INTRODUCTION

Mitochondria play critical roles in maintaining cellular homeostasis, generating ATP, buffering calcium, but sometimes at the cost of generating reactive oxygen species (ROS) [1]. Mitochondria are sophisticated organelles that require the precise coordination of protein homeostasis with resident proteins transcribed, translated, and imported from the nucleus, but also proteins coded within the mitochondrial genome. To maintain a healthy lifespan, organisms must prevent the accumulation of protein aggregation by targeting misfolded proteins for proteasomal degradation or removal by autophagy [2]. The inability to maintain homeostatic function of the mitochondrial proteome can lead to organelle disruption and even loss of cellular functions [3].

Given its nickname as the powerhouse of the cell, the electron transport chain (ETC) is required for efficient oxidative phosphorylation (OXPHOS), supporting ATP synthesis *en masse* [4]. However, the ETC and complex I that passes electrons from NADH to ubiquinone is a major generator of ROS. Mitochondria contain a protective arsenal of enzymes to minimize ROS, including the capacity to dismutase superoxide anions to hydrogen peroxide (H_2_O_2_) and catalases which can further catabolize H_2_O_2_ to water and molecular oxygen with the assistance of glutathione antioxidant systems [5]. In the absence of these protective measures, OXPHOS deficiency promotes mitochondrial dysfunction and cell damage that drives several age-related pathologies (e.g., dementia, sarcopenia, and insulin resistance) [6-9] .

WDR23, as a part of the cellular proteostasis machinery, regulates cytoprotection in metabolic tissues, but the functions of WDR23 in the brain are not documented. As part of the CUL4-DDB1 E3 ubiquitin ligase, WDR23 is an adapter for substrates that are targeted for degradation. Several specific substrates have been identified, including SKN-1/NFE2l2 [10], GEN1 [11], SLBP [12], and p21 [13]. Although a mitochondrial resident protein has yet to be established as a direct target of WDR23, several of the established direct targets play critical roles in mitochondrial homeostasis.

Mitochondria contain thousands of proteins of which only 13 are encoded by the mitochondrial genome with most mitochondrial resident proteins being nuclear-encoded, translated in the cytoplasm, and imported to their correct subcompartment within the mitochondria by translocases of the outer mitochondrial membrane (TOM) and translocases of the mitochondrial inner membrane (TIM) [14]. As such, mitochondrial biogenesis and maintenance requires a coordinated effort across multiple cellular compartments for proper function. Nuclear respiratory factor 1/2 (NRF1/2) transcribes nuclear genes that encode ETC proteins, transcription factor A (TFAM), and other proteins critical for mitochondrial biogenesis [15].

In this work we study the effects of genetic ablation of *Wdr23* in the hippocampus of the murine brain, which importantly alters steady state levels of mitochondrial proteins involved in OXPHOS and stress adaptation. These alterations could play critical roles in the homeostatic regulation of aging and aging-associated degradation of mitochondrial functions. Our work provides a new molecular player and an important biomarker of mitochondria function that could be modulated to ensure healthy aging.

## MATERIALS AND METHODS

### Animals

All animal protocols were approved by the Institutional Animal Care and Use Committee (IACUC) of the University of Southern California and all the procedures were conducted in compliance with institutional guidelines and protocols.

*Wdr23* knock-out (*Wdr23*KO) mice were generated by Wellcome Trust Sanger Institute [16, 17]. *Wdr23KO* animals were subsequently backcrossed nine times into our C57BL/6J (WT) strain from the Jackson laboratory. Heterozygous (*Wdr23 +/-*) *dams* and *sires* were then mated to generate *Wdr23+/*+ and *Wdr23-/-* animals that were maintained as WT and KO, respectively. Male mice (n=4-6/group) were kept in a 12:12 h light-dark cycle, constant temperature, and humidity room. All animals were allowed *ad libitum* access to water and food.

### Rodent diets

Mice were fed ad lib with irradiated D12450K rodent chow (Research Diets), containing 16.1kJ/g of digestible energy (animal-based protein 3.22kJ/g, carbohydrate 11.27kJ/g, fat 1.61 kJ/g).

### RNA extraction and RNA sequencing

The hippocampus tissue of 44 weeks old mice was collected and lysed in RNeasy Lipid Tissue mini kit reagent (QIAGEN). RNA was extracted according to the manufacturer’s protocol. Qubit RNA BR Assay Kit was used to determine RNA concentration. Isolated RNA was sent to Novogene for library preparation and deep sequencing in biological triplicate. The read counts were used for differential expression (DE) analysis by using the R package DEseq2 (R version 3.5.2). Differentiated expressed genes were analyzed using p value <0.05 and fold change >1.5 as cutoff.

### Western blot analysis

The hippocampus tissue of 8-14 weeks old mice was collected for WB analysis. Whole cell lysates were prepared in M-PER buffer (1x Mammalian Protein Extraction Reagent (Thermo Scientific), 0.1% Halt Protease & Phosphatase inhibitor (Thermo Scientific) according to the manufacturer’s protocol. Total protein concentrations were quantified by Bradford assay (Sigma). An equal amount of protein (20 µg) was separate on 4%-12% bis-tris polyacrylamide gel (Invitrogen,) in MOPS running buffer (Invitrogen) and then transferred to nitrocellulose membranes (GE Healthcare Life science). After blocking for 1 h with 3% BSA in PBST (PBS, 0.1% Tween 20), the membranes were subjected to immunoblot analysis. Antibodies used include: IDE (Millipore sigma, 1:5000), HO-1 (Abcam, 1:2000), TOMM20 (Abcam, 1:50000), AIF (Abcam, 1:1000), PGC1β (Abcam, 1:5000), TIMM17A/TIM17 (Abcam, 1:1000), VDAC1/porin+VDAC2 (Abcam, 1:5000), GCLC (Cell signaling, 1:1000), RXFP3 (Novus Biologicals, 1:1000), GSTP1 (Novus Biologicals, 1:1000), RLN3 (Novus Biologicals, 1:1000), β-actin (Millipore Sigma, 1:10000) and HRP-conjugated secondary antibodies (Thermo Fisher, 1:10,000). Specific protein bands were visualized and evaluated using FluorChem HD2 (ProteinSimple).

### Statistical Analysis

All experiments were performed at least in triplicate. Data are presented as mean ± SEM. Data handling and statistical processing were performed using GraphPad Prism 8.0. Comparisons between two groups were done using unpaired Student’s t-test. Differences were considered significant at the p ≤ 0.05 level. *p<.05, **p<.01, ***p<.001, ****p<.0001, compared to C57BL/6J (WT) control.

## RESULTS

### *Wdr23KO* animals increase expression of NRF1-regulated transcripts in the hippocampus

The brain is under energetic constraints that must continuously balance basal cellular function, repair damage, and monitor molecular turnover throughout the lifespan. We first measured the transcriptional response to loss of *Wdr23* by performing RNA-sequencing on dissected hippocampus from *Wdr23KO* mice and compared to samples from age-matched WT animals. 46 genes displayed a significant difference in expression (**Table 1**). Among the classes of genes most highly upregulated include proteins with roles in neuronal function (e.g., synuclein, synaptogamin, potassium voltage-gated channels), cellular signaling (e.g., G protein coupled receptors, nicotinic cholinergic receptors), and metabolism (e.g., phospholipase C, aldehyde dehydrogenase family 1, prostaglandin synthase).

**Table 1.**
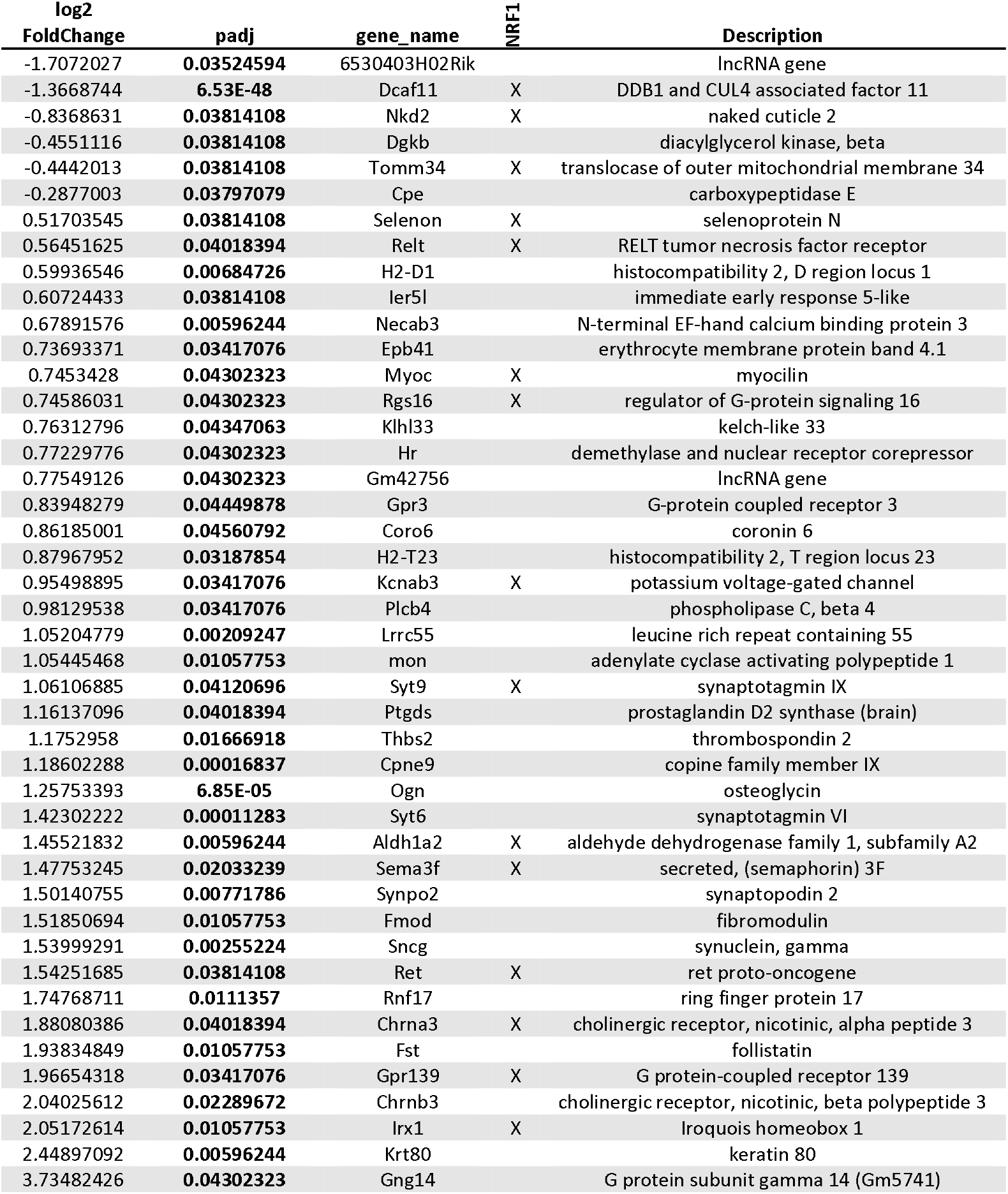
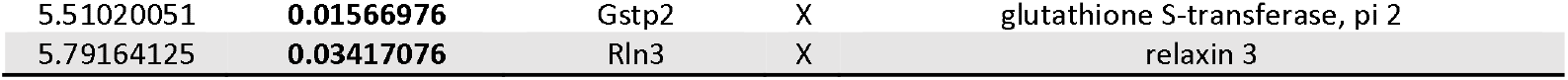
RNAseq of hippocampus (include annotation of NRF binding sites)

We also noted a significant reduction in the expression of translocase of the outer mitochondrial membrane 34 (Tomm34) protein, which plays a role in the import of nuclear-coded mitochondrial proteins across the mitochondrial outer membrane. Moreover, 37% of the dysregulated genes have binding sites in the promoter region of the gene for nuclear respiratory factor 1 (NRF1) [18, 19]; a key mediator of genes involved in oxidative phosphorylation (OXPHOS) and mitochondrial biogenesis [20, 21]. Taken together, the transcriptional signature suggests a change in mitochondria in the brain of animals lacking *Wdr23*.

### Loss of Wdr23 results in OXPHOS complex imbalance in the hippocampus

The function of the ETC requires the coordination of protein import and assembly of nuclear and mitochondrial subunits, where imbalances in this system can lead to defects that drive the generation of reactive oxygen species (ROS) and cellular dysfunction. Based on the established role that NRF1 plays in regulating oxidative phosphorylation, we first looked for changes in the mitochondrial ETC (**Figure 1A**). We measured the abundance of the five ETC complexes by Western blot with antibodies that specifically recognize NADH dehydrogenase [ubiquinone] 1 beta subcomplex subunit 8 (NDUFB8) in complex I, succinate dehydrogenase [ubiquinone] iron-sulfur subunit (SDHB) in complex II, cytochrome b-c1 complex subunit 2, mitochondrial (UQCRC2) in complex III, mitochondrially encoded cytochrome c oxidase I (MTCO1) in complex IV, and ATP synthase F1 subunit alpha (ATP5A) in complex V (**Figure 1B-G**). Rather than a change in all ETC complex components, we measured a significant increase in the subunits for complex I (NDUFB8, **Figure 1C**) and complex III (UQRC2, **Figure 1E**), but not complex II (SDHB, **Figure 1D**), complex IV (MTCO1, **Figure 1F**), or complex V (ATP5A, **Figure 1G**).

**Figure 1.**
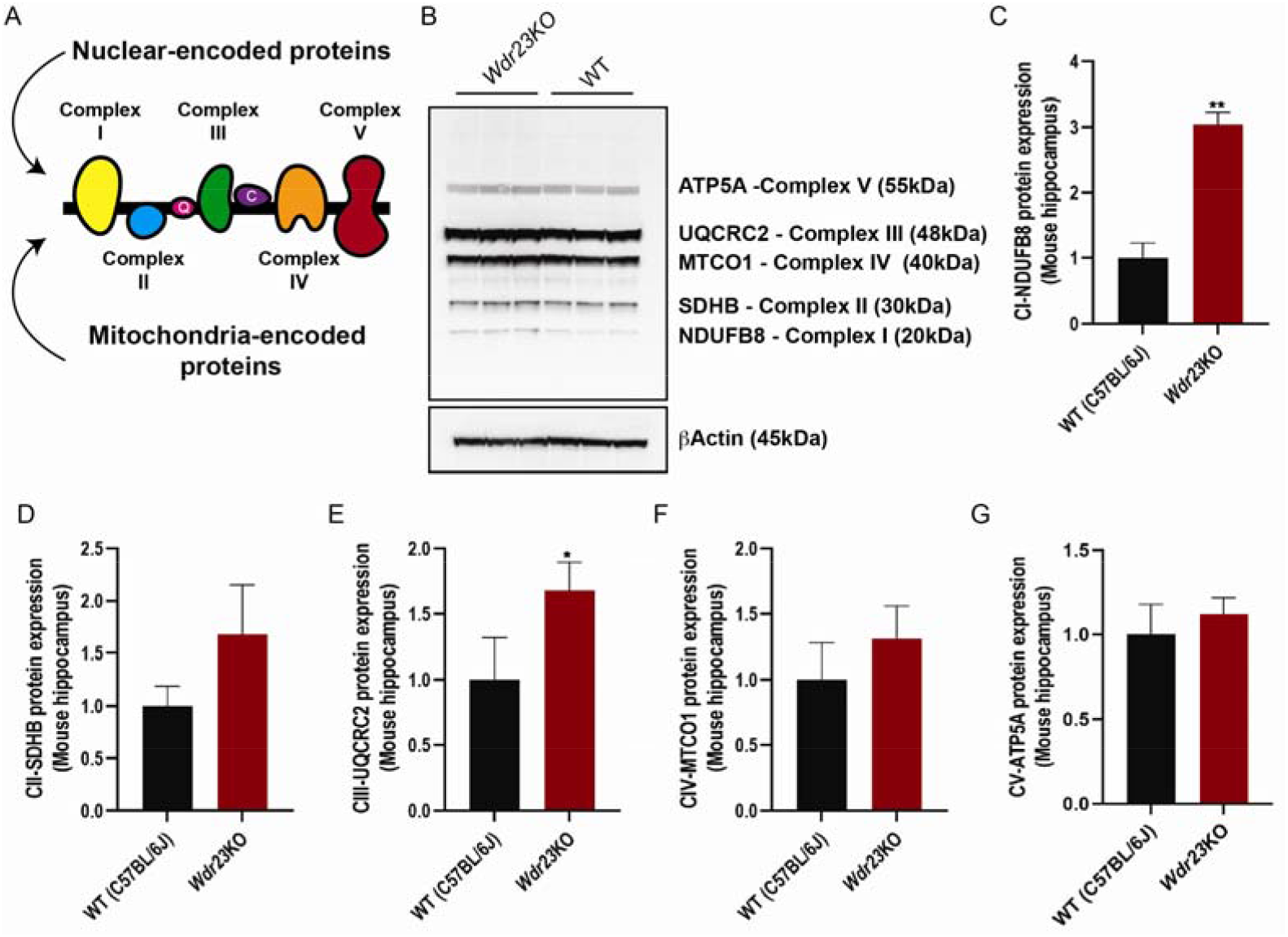
Mitochondrial OXPHOS complex I and III are increased in the brain of *Wdr23KO* mice. (A) Cartoon model of ETC complexes. (B) Western blot analysis of total protein from hippocampal homogenates probed for subunits of the mitochondrial electron transport chain complexes I-V and quantification of differences between WT and *Wdr23KO* samples, normalized against *β*-actin, for (C) NDUFB8 of complex I, (D) SDHB of complex II, (E) UQCRC2 of complex III, (F) MTCO1 of complex IV, and (G) ATP5A of complex V.

### *Wdr23KO* displays increased expression of stress signaling proteins

We next examined the steady state level of several stress response pathways in homogenates of dissected hippocampi from WT and *Wdr23KO* animals (**Figure 2A**). Our previous studies of liver tissue from the *Wdr23KO* mouse revealed activation of NFE2l2/NRF2 pathway due to the negative regulation of this transcription factor by the Cul4-WDR23 ligase and the ubiquitin proteasome system [10, 11]. Unsurprisingly, we noted an increase of log2fold change of 5.5 in the transcript levels of the established NFE2l2/NRF2 target gene Glutathione s-transferase,pi 2 (*Gstp2*) (**Table 1**). We confirmed a significant increase in the steady state level of GSTP proteins with an antibody that recognizes all three orthologous GSTP isoforms (**Figure 2B**). We next measured the expression of mitochondrial apoptosis inducing factor (AIF) that helps mediate the biogenesis and maintenance of complex I [22]. We noted a modest, but significant increase in the steady state levels of AIF (**Figure 2C**) and the voltage-dependent anion channel (VDAC1) of the outer mitochondrial membrane (**Figure 2D**). Although we measured a clear enrichment for NRF1 targets, we could not detect a difference in the abundance of the upstream regulator of NRF1 activation, PGC1*α*/*β* [23] (**Figure 2E**). Lastly, we measured the steady state level of the TOMM20, a member of the translocase of the mitochondrial outer membrane, which was significantly reduced (**Figure 2F**), similar to transcript levels of the TOMM complex partner Tomm34 (**Table 1**). Taken together, these results demonstrate that in addition to changes in the abundance of distinct OXPHOS complex proteins (**Figure 1**), several mitochondrial resident proteins with roles in stress responses and organization are altered in the hippocampus of *Wdr23KO* mice.

**Figure 2.**
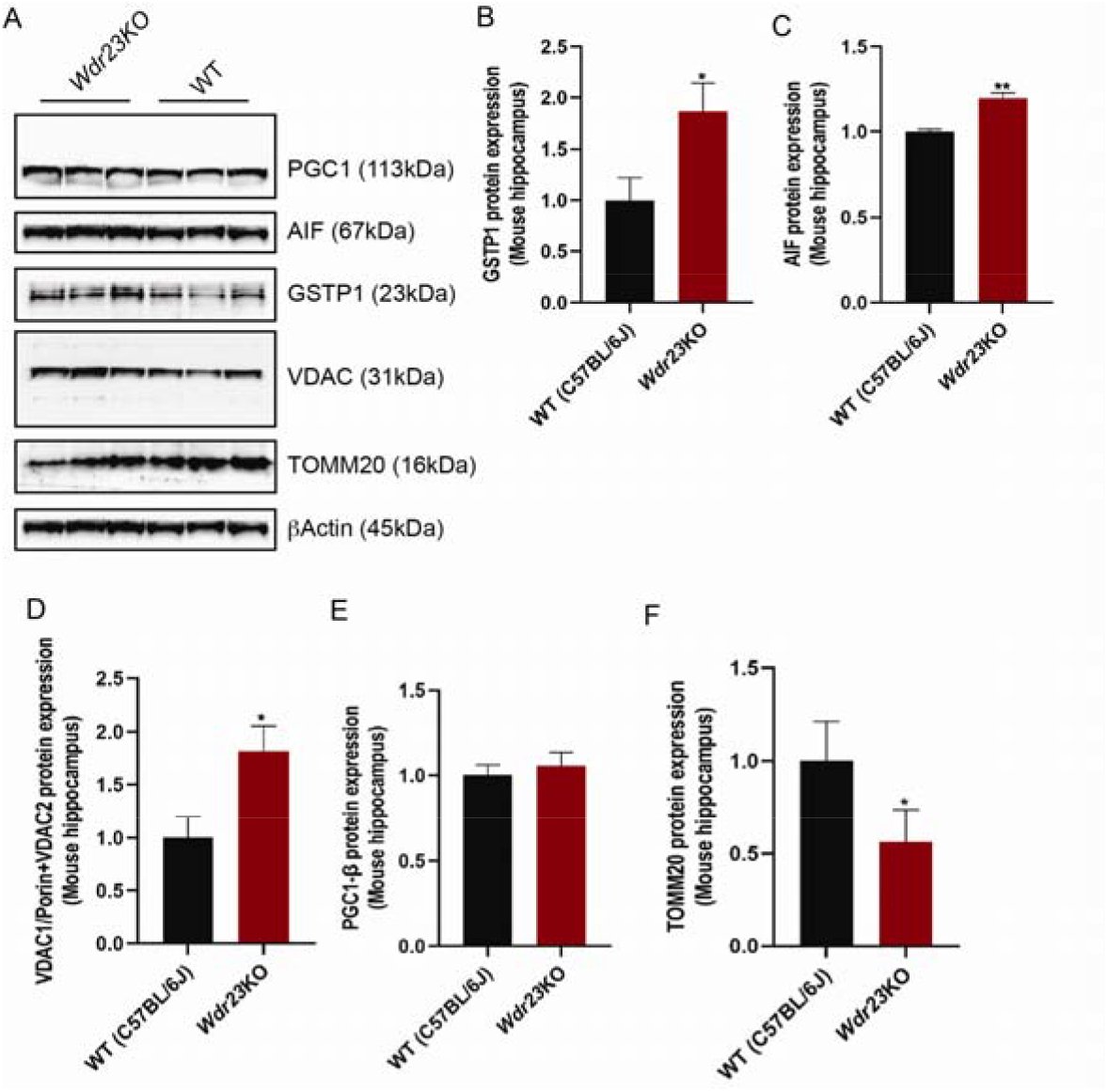
Loss of *Wdr23* proteostasis increases mitochondrial stress factors. (A) Western blot analysis of total protein from hippocampal homogenates probed for proteins with roles in stress adaptation and homeostasis. Differences between WT and *Wdr23KO* samples were quantified and normalized against *β*-actin, for (B) GSTP1, (C) AIF, (D) VDAC1/2, (E) PGC1*β*, and (F) TOMM20. Comparisons analyzed by t-test; *p<0.05, **p<0.01

Similar to oxidative stress pathway genes that are activated in *Wdr23KO* mice [10, 11], our recent work revealed an increase in the expression of the insulin degrading enzyme (IDE) in the liver of animals lacking WDR23 proteostasis [24]; noting that IDE can localize to the mitochondria [25, 26]. Surprisingly, the most highly expressed gene in the hippocampus of *Wdr23KO* mice, relative to WT, was the relaxin-3 locus, which was upregulated log2fold change of 5.7 (**Table 1**), which has established roles in cellular responses to oxidative stress [27]. As such, we measured the steady state abundance of these two proteins in our hippocampus homogenates from *Wdr23KO* and WT animals (**Figure 3A**). We noted a ∼3.5-fold increase in the abundance of IDE (**Figure 3B**) and a >1.5-fold increase in RLN3 (**Figure 3C**). Notably, we could not detect a change in the steady state level of the Relaxin Family Peptide Receptor 3 (RXFP3, **Figure 3D**). Taken together, these data suggest that in addition to changes in the abundance of essential mitochondrial resident proteins in the brain tissue of *Wdr23KO* animals, signaling molecules, and metabolic regulators are significantly altered in response to loss of *Wdr23*.

**Figure 3.**
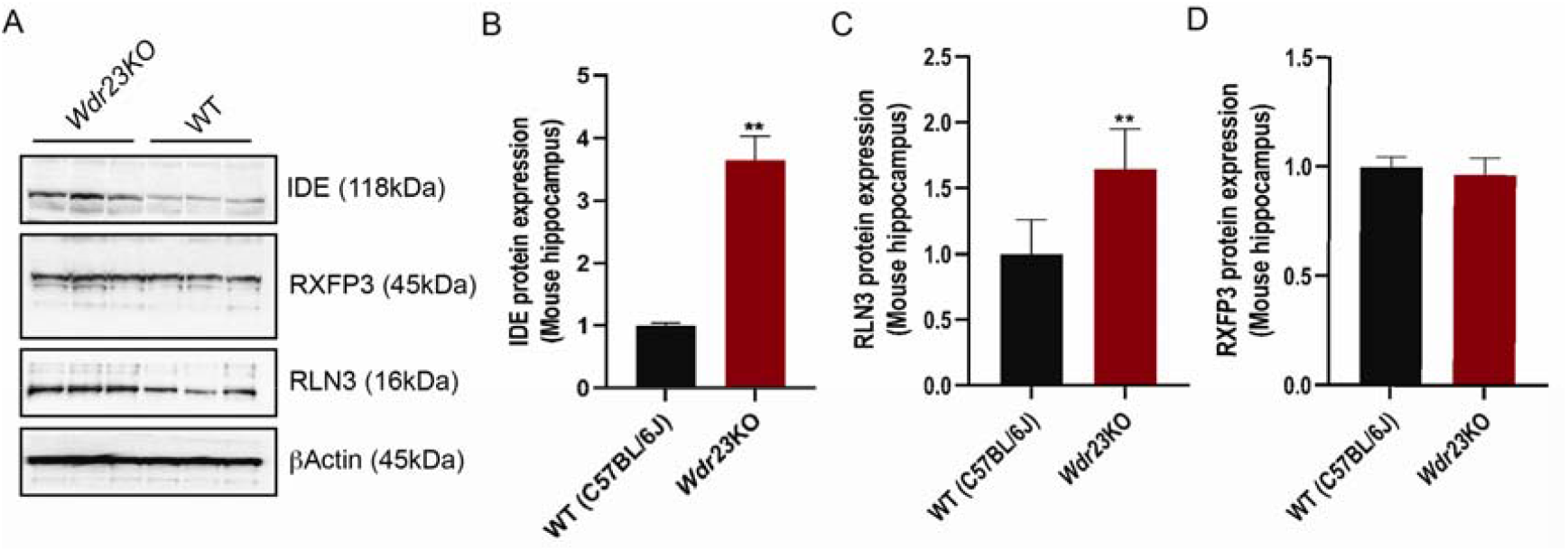
Wdr23KO mice display increased IDE and RLN3 expression in the hippocampus. (A) Western blot analysis of total protein from hippocampal homogenates probed for proteins with roles in metabolic signaling. Differences between WT and *Wdr23KO* samples were quantified and normalized against *β*-actin for (B) IDE, (C) RLN3, and (D) RXFP3. Comparisons analyzed by t-test; **p<0.01

## DISCUSSION

Loss of coordination between the subunits within each complex of the electron transport chain and across the complexes themselves can lead to reduced mitochondrial function (e.g., reduced ATP and increased ROS). To correct this imbalance, the mitochondria and nucleus utilize sophisticated signaling pathways to communicate, which if unsuccessful, can lead to additive effects of age-associated damage to the organelle and the cell. Our understanding of the cause and consequence of loss of proteostasis and aging is complicated by the fact that endogenous and exogenous stressors accumulate with age and can promote the aggregation of misfolded proteins when cellular chaperones are unable to correct the defect which drives age-related pathologies. Moreover, with age, the activity of these chaperones is reduced resulting in the further imbalance of proteostasis. Understanding the homeostatic mechanisms that protect mitochondrial health through the integration of nuclear and mitochondrial signaling pathways to support metabolic plasticity and stress responses in the nervous system is of great importance. We identify *Wdr23*, a member of the Cul4-DDB1 E3 ubiquitin ligase complex, as a new molecule with the capacity to modulate the steady state abundance of mitochondrial resident proteins. The identification of new biomarkers that maintain homeostasis is of great importance and provides new molecular approaches for genetic, molecular, and pharmacological manipulation of physiological systems.

We identify three proteins with established roles in regulating the complexes of the mitochondrial OXPHOS system that are upregulated in the absence of *Wdr23*. AIF is essential for oxidative phosphorylation and has been linked to the stabilization of complex I and III, both of which display increased abundance in *Wdr23KO* brain [28]. AIF itself can also translocate to the cytoplasm and eventually the nucleus where it can regulate cell death [22, 28], however the abundance of AIF was only modestly increased in *Wdr23* hippocampi and perhaps not to the level necessary to drive apoptosis. This finding is correlated with the increased abundance of VDAC1/2 in hippocampal homogenates from *Wdr23KO* animals. The accumulation of VDAC1 can increase the mitochondrial permeability transition pore (mPTP) opening [29, 30] under mitochondrial stress conditions, but whether the increase in VDAC1 is a response or result of *Wdr23* loss remains to be elucidated.

Second, IDE, which can localize to the mitochondria [25], and beyond its role in regulating the levels of insulin, can interact with multiple mitochondrial ribosomal proteins in addition to proteins that facilitate the synthesis and assembly of mitochondrial complex I and IV [26]. Both complex I and IV are increased in the *Wdr23KO* brain albeit the latter failed to reach significance, but the trend for increased MTCO1 of complex IV was observed. Loss of complex I integrity, and the resulting imbalance of metabolism and ROS production has been implicated in metabolic diseases including diabetes and neurological conditions such as mood disorders and Parkinson’s disease [6].

Third, the most differentially expressed transcript in the hippocampus of *Wdr23KO* mice, as compared to WT, was *Rln3*, which encodes an insulin-like peptide linked to age-related neurological conditions including memory dysfunction, appetite control, and depression [27]. In addition, relaxin peptides have been used pharmacologically to provide protection from ischemia, which is thought to occur through the activation of mitochondrial respiratory pathways [31]. Rodent and non-human primate models suggest relaxin-3 signals through the G-protein coupled receptor, RXFP3, which has a broad range of effects on neuroendocrine functions associated with stress responses [32]. However, unlike the increase measured in RLN3 signaling peptide, we did not observe a similar increase in RXFP3, suggesting the loss of *Wdr23* genetically mimics the pharmacological use of RLN3 *in vivo*.

Taken together our study documents an increase in the abundance of specific mitochondrial resident proteins in the hippocampus of animals harboring a genetic mutation in *Wdr23*. Moreover, the loss of *Wdr23* results in significant increases in AIF, IDE, and RLN3, all of which have established roles in regulating ETC complexes and OXPHOS activity. Future work to define the age-related impact of loss of *Wdr23* on these molecules will be of great interest.

## ACKNOWLEDGEMENTS

We thank M. Donoghue, M. Lynn and S. Ledgerwood for technical assistance. We thank C. Ramos and Dr. N. Stuhr for critical reading of the manuscript. This work was funded by the NIH RF1AG063947 to SPC and a Glenn Foundation for Medical Research Postdoctoral Fellowship in Aging Research from the American Federation for Aging Research to CD.

## Author contributions

Conceptualization: SPC; Methodology: CD, RI, and SPC; Investigation: CD, RI, and SPC; Visualization: CD and SPC; Supervision: SPC; Writing (original draft): SPC; Writing (reviewing & editing): CD and SPC

## Data and materials availability

All data are available in the main text or the supplementary materials.

